# Seize the Data: A User-Friendly GUI for High-Resolution Analysis of Seizure Dynamics in HD-MEA Recordings

**DOI:** 10.64898/2026.01.07.698219

**Authors:** Melissa L. Blotter, Jacob Cahoon, Jacob H. Norby, Katelyn C. Forbes, Logan A. Stephens, Grace A. Schmutz, Micah R. Shepherd, R. Ryley Parrish

**Affiliations:** Neuroscience Center, Brigham Young University; Department of Cell Biology and Physiology, Brigham Young University; Department of Computer Science, Brigham Young University; Department of Physics and Astronomy, Brigham Young University

**Keywords:** Electrophysiology, Neuroscience Tools, Microelectrode Array, Seizure Dynamics, Seizure Analysis Tool

## Abstract

High-density multi-electrode arrays (HD-MEAs) generate large, complex datasets that are challenging to efficiently manage and analyze with existing tools, especially in open-source environments. To address this, we developed the BYU Seizure and Analytics Tool (YSA), an open-source graphical user interface (GUI) built in Python and C++ for efficient analysis and visualization of HD-MEA recordings. The YSA features raster plots, automated discharge detection and tracking, downsampling, playback, and export functions, enabling streamlined workflows for large-scale neural data. We demonstrate the utility of the tool in the context of seizure and status epilepticus (SE)-like activity, highlighting how the YSA facilitates rapid exploration of the spatiotemporal dynamics in brain networks. This platform provides an accessible and practical solution for HD-MEA data analysis, supporting a range of neuroscience applications.

**Significance Statement:** High-density multi-electrode arrays (MEAs) generate rich spatiotemporal datasets ideal for studying complex brain dynamics such as seizure activity. However, the size and complexity of these data often pose challenges for efficient analysis and interpretation. We present the BYU Seizure and Analytics Tool (YSA), an open-source graphical interface for intuitive visualization and exploration of MEA data. YSA enables users to navigate activity across the entire brain slice, identify relevant patterns, and easily export subsets for targeted analyses such as individual discharges. By making high-density seizure data more accessible and actionable, YSA streamlines analysis workflows and supports deeper insights into the spatiotemporal dynamics underlying seizure initiation and propagation in models of status epilepticus, pharmacoresistant epilepsy, and broader neural activity.

## Introduction

The ability to monitor neural activity across thousands of electrodes simultaneously has transformed the landscape of neural recording, thanks to advances in high-density multi-electrode array (HD-MEA) systems (Blotter et al. 2024; Maccione et al. 2010; van Vliet et al. 2007). These systems are increasingly used in neuroscience and other fields to study network-level phenomena with exceptional spatial and temporal precision (Liu et al. 2012). However, the volume and complexity of data generated by HD-MEAs present major challenges for data handling, analysis, and visualization, particularly for researchers lacking access to proprietary or specialized software(Muller et al. 2015; Obien et al. 2014).

To address this need, we developed the BYU Seizure and Analytics Tool (YSA), an open-source graphical user interface (GUI) built in Python and C++. YSA is designed specifically to streamline the analysis of HD-MEA recordings, integrating essential features such as customizable raster plots, automated discharge detection, real-time playback, and easy data export into a single, user-friendly platform. Its flexible design enables researchers to quickly explore spatiotemporal patterns in large-scale electrophysiological data in a fully customizable and open-access environment, while also remaining accessible to users with limited programming experience.

Several open-source tools for MEA analysis are currently available, including platforms like SpyKING CIRCUS (Yger et al. 2018), MEAnalyzer (Dastgheyb, Yoo, and Haughey 2020), and NeuroScope (Hazan, Zugaro, and Buzsaki 2006), and the Xenon LFP Analysis Platform (Mahadevan, Codadu, and Parrish 2022). While these tools offer powerful capabilities, particularly for spike sorting, burst detection, and traditional LFP analysis, many are optimized for single-unit or burst-level dynamics and are less suited for visualizing and tracking complex spatiotemporal seizure activity. For example, the Xenon LFP Analysis Platform provides basic signal viewing and spectral features but lacks the customizable discharge-level tracking, interactive visualization, and modular flexibility needed for modern HD-MEA seizure analysis. Additionally, many of these platforms are not easily adaptable to evolving recording technologies or require substantial scripting experience. In contrast, YSA offers a real-time, user-friendly interface that enables detailed per-discharge analysis, synchronized raster and trace visualization, seamless data export, and compatibility with current and future MEA systems. Its modular, open-source design supports easy integration of user-defined routines, making it uniquely suited for high-throughput and translational seizure research.

To showcase the analytical capabilities of YSA in a real-world context, we applied it to *ex vivo* recordings from an acute seizure model. This seizure model presents complex, dynamic patterns of seizure-like and status epilepticus (SE)-like activity (Reddy and Kuruba 2013; Voss and Sleigh 2010; Walther et al. 1986; Codadu et al. 2019; Codadu, Parrish, and Trevelyan 2019), making it an ideal test case for evaluating the tool’s ability to detect, visualize, and quantify large-scale network phenomena. While our study is grounded in epilepsy research, the tool is broadly applicable to any experimental paradigm involving HD-MEA data. The seizure model used here, including sustained SE-like activity that mimics pharmacoresistant seizure states, offers a compelling demonstration of YSA’s ability to dissect complex network dynamics. User-specific routines can be easily added to the GUI for custom applications. YSA offers a robust, extensible solution for high-throughput neural data analysis and contributes to the growing ecosystem of open-source tools in neuroscience.

## Materials and Methods

### Animals and brain slice preparation

Acute horizontal brain slices were prepared from male and female C57BL/6 mice aged postnatal day (P)17 to P35. Mice were anesthetized with isoflurane and euthanized via decapitation, in accordance with approved institutional animal care protocols. Brains were rapidly extracted and immersed in a cold cutting solution containing (in mM): 126 NaCl, 3.5 KCl, 1.26 NaH₂PO₄, 26 NaHCO₃, 10 glucose (C₆H₁₂O₆), and 3 MgCl₂. Brains were sectioned into 350 µm-thick horizontal slices using a Leica VT1200S vibratome (Precisionary Instruments, Ashland, Massachusetts, USA). Slices were sub-dissected to isolate the neocortex, entorhinal cortex, hippocampus, and subiculum (Figure 1A).

**Figure 1:**
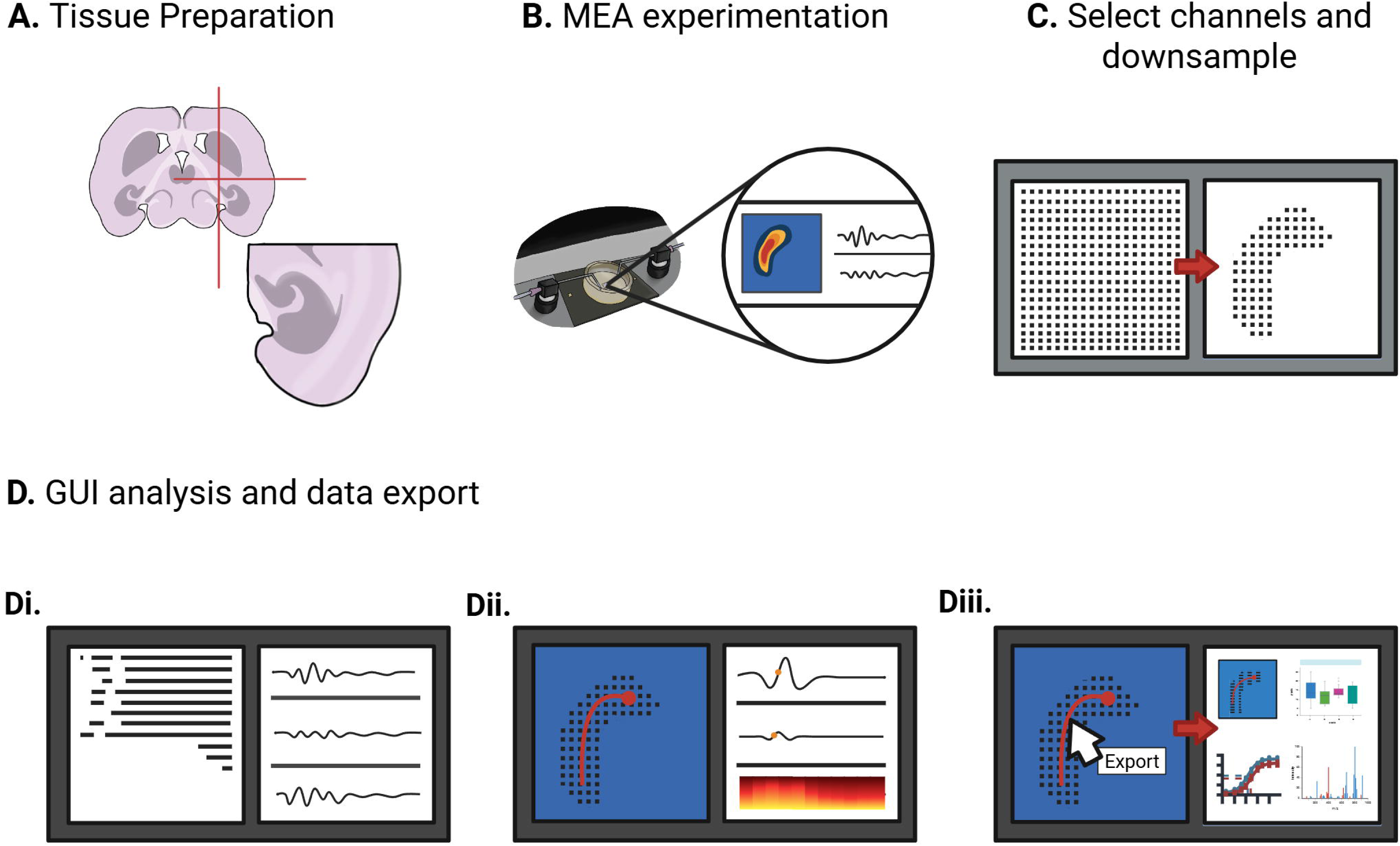
Workflow of experimentation and analysis. **A)** Cartoon of an acute mouse horizontal brain slice, that includes the hippocampus, neocortex, and entorhinal cortex. **B)** Brain slice placed onto the MEA where seizure activity is induced and recorded. **C)** Raw recordings are downsampled spatially and temporally to facilitate fast analysis. **D)** The processed data are loaded into the YSA, which **Di)** generates raster plots, **Dii)** enables visualization of individual discharges and spectrograms, and **Diii)** allows for easy export of data for downstream analysis.

Following dissection, slices were transferred to a holding chamber containing artificial cerebrospinal fluid (aCSF) composed of (in mM): 126 NaCl, 3.5 KCl, 1.26 NaH₂PO₄, 26 NaHCO₃, 10 glucose, 1 MgCl₂, and 2 CaCl₂ for a one-hour incubation. All solutions were continuously bubbled with carbogen (95% O₂ / 5% CO₂) to maintain proper oxygenation and pH.

### Electrophysiological recordings on HD-MEA

After incubation, slices were placed on a 3Brain Accura high-density multi-electrode array (HD-MEA) chip containing 4,096 electrodes spaced 60 µm apart, arranged in a 3.8 × 3.8 mm grid (Figure 1B). Recordings were performed using the 3Brain BioCAM DupleX system (3Brain, Paffikon, Switzerland). Prior to induction of epileptiform activity, slices were perfused with standard aCSF (composition described above) for 10 minutes to allow for recovery and equilibration. Slices were then perfused in a low Mg²⁺ artificial cerebrospinal fluid (aCSF) solution composed of (in mM): 126 NaCl, 3.5 KCl, 1.26 NaH₂PO₄, 26 NaHCO₃, 10 glucose, and 2 CaCl₂, to induce epileptiform activity. The solution was continuously perfused at an inflow rate of 5.0 mL/min and an outflow rate of 7.0 mL/min and maintained at physiological temperature (35–37 °C) using an in-line heating system (Warner Instruments, Holliston, Massachusetts, USA). Each recording session lasted up to 3 hours and was conducted as previously described (Blotter et al. 2024).

Images of each brain slice on the 3Brain Accura chip were captured using a Dino-Lite Edge digital microscope to ensure precise localization of recording channels relative to the targeted brain regions (Dunwell Tech., Inc. Dino-Lite US). Data were sampled at 2,000 Hz with a 1.0 Hz high-pass filter applied and no low-pass filter.

### Data preprocessing

Recorded data were downsampled both spatially and temporally using custom code which is publicly available and designed for the 3Brain recordings used in this study (Lab 2025). The recordings were downsampled spatially by manually selecting channels on the array that correspond to the location of the slice, eliminating data from irrelevant channels (Figure 1C). Temporally, the recordings were downsampled from 2000 Hz to 100 Hz to remove extraneous frequencies. While temporal down-sampling is not necessary, it will reduce the CPU time required to load the data into YSA and perform analysis, without losing any integrity of results. Data conversion to an HDF5 file format is required for YSA use; example conversion code and the necessary data structure for YSA-compatible HDF5 files are available on the readthedocs guide (Lab 2025).

### Custom GUI for seizure detection

#### Overview and usage

The downsampled data were imported into YSA (Figure 1D & Figure 2A). Upon loading, users are presented with a control panel for importing data, adjusting parameters, and navigating the interface (Figure 2A). A sidebar menu (turquoise) allows access to various tabs, including a visualization of the MEA grid with active electrode positions (Figure 2A). Users can highlight up to four individual channels to display the signals with quick zoom-in and -out and pan options. Switching to the raster plot tab displays spike activity across all electrodes over time, with detected self-limiting seizure-like events (SLSLEs) (light blue) and SE events (yellow) overlaid automatically (Figure 2B). Naturally, detection routines can be updated (with only small amounts of coding) to fit the user’s specific needs. Raster plots can also be displayed alongside trace plots for any selected channels, allowing close inspection of raw signals during identified events (Figure 2C). A minimap panel (pink) provides an overview of the entire recording timeline, indicating the currently displayed window.

**Figure 2:**
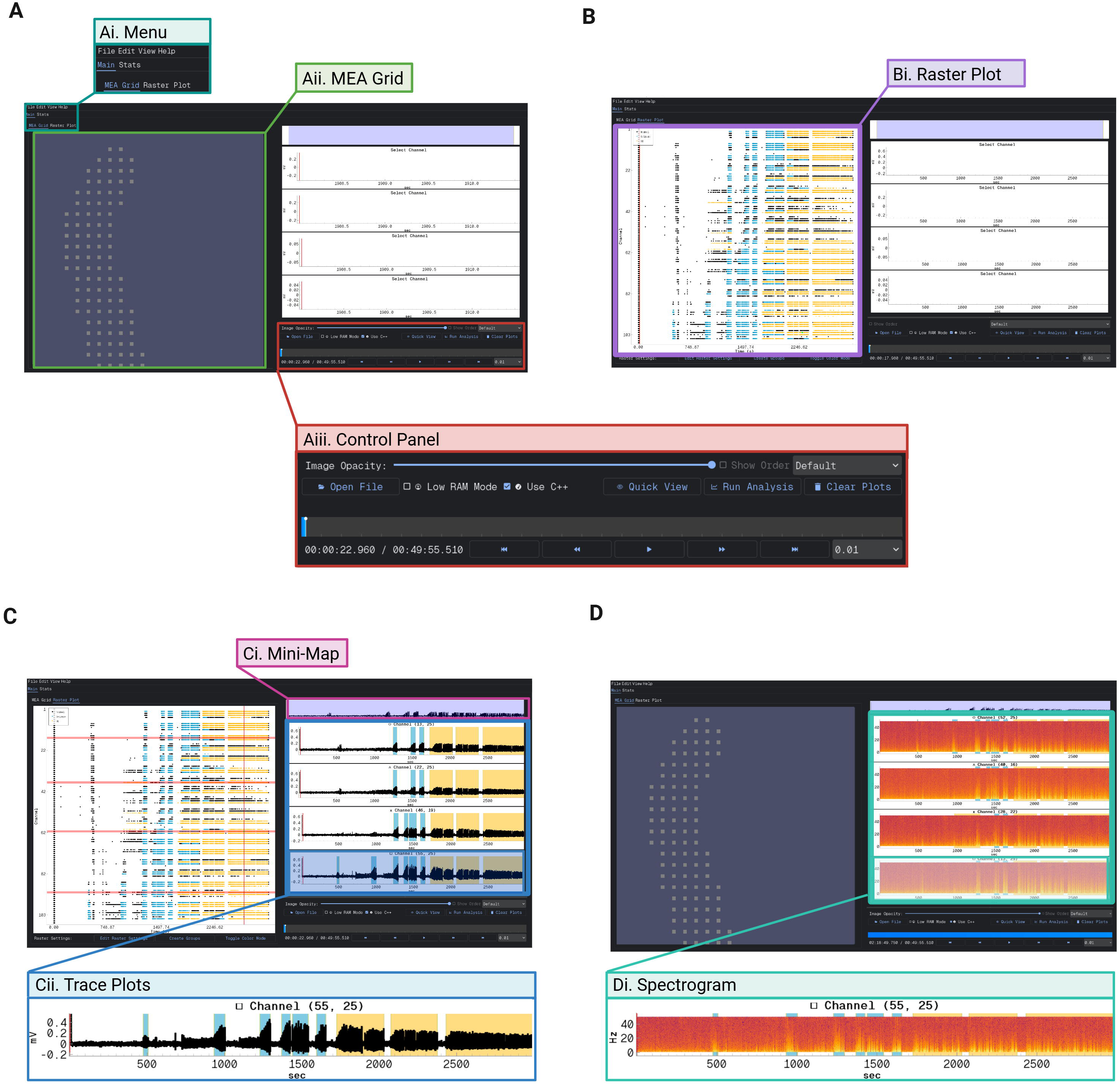
YSA Overview. **A)** Main interface of the YSA tool. **Ai)** The full GUI view with the menu panel highlighted (turquoise box), which allows users to navigate between tabs. **Aii)** The MEA Grid tab (green box) showing the electrode layout and signal overview across the array. **Aiii)** The control panel (red box), which includes options for uploading recordings, adjusting playback speed, and navigating the dataset. **B)** Raster plot view. **Bi)** The Raster tab (purple box) displays network-wide spike activity, with automated annotations for seizure-like events (light blue) and status epilepticus (SE)-like activity (yellow). **C)** Trace plot view. **Ci)** The minimap timeline (pink box) provides a compressed view of the full recording for navigation. **Cii)** Individual trace plots (blue box) display voltage traces from selected electrodes. One trace is enlarged to show axis labels and temporal detail. **D)** Spectrogram view. **Di)** Example spectrograms from individual electrodes (light green box), showing frequency content over time. A combined trace-spectrogram view is enlarged to display axes and resolution.

In addition, users can view time-frequency representations of activity via per-electrode spectrograms (Figure 2D), enabling frequency-domain inspection of discharges and their evolution across the array. Spectrogram parameters can be modified within the edit settings menu.

#### Seizure detection algorithm

YSA can be used to identify regions that exhibit self-limiting seizure-like events (SLSLEs) and SE-like activity using a seizure detection algorithm. This algorithm works in three steps: First, it identifies a reference or baseline section of a 2-minute period with no significant activity spikes. Next, it identifies voltage outliers that deviate by 3 standard deviations (SD) from the baseline. For each of these, the algorithm checks whether the variance in that window also deviates by 3 SD from the reference variance. The resulting set of points are defined as discharges. Discharges occurring within 2 seconds of each other are grouped together in the same event. These events are then categorized by their duration: those lasting longer than 10 seconds are classified as SLSLEs, those shorter than 10 seconds are discarded, and those lasting longer than 5 minutes are classified as SE (Dulla et al. 2018). Improved detection routines or other custom algorithms can be easily added to the source code to replace the default algorithm.

### Tracking and mapping discharges

The YSA GUI includes a peak detection module to identify all discharge peaks across electrodes. With user set parameters for minimum peak threshold (in SD deviation from the mean), minimum distance between peaks (in samples), and signal to noise ratio, peaks are identified in each channel. For each peak, the preceding time window is analyzed to locate the point with the largest voltage difference, marking this as the discharge onset. This approach ensures alignment between detected discharges and their representation on the false color map. Once peaks are identified, YSA tracks their propagation across the slice using the DBSCAN algorithm (from the scikit-learn python package, which also has tunable parameters in the settings menu) to place the center of the discharge at each time point as the discharge travels. Additionally, it allows users to mark and monitor the electrodes involved in discharge initiation, enabling analysis of spatial discharge patterns and timing intervals between events.

### Statistical Analyses

Statistical analyses of data were performed on .csv outputs from the YSA using custom MATLAB code and automated where possible. Data of two groups were analyzed with two-tailed t-tests for all plots. Figures were made in MATLAB (The MathWorks, Natick, MA, USA) and in BioRender (BioRender, Toronto, Ontario, Canada).

### Software Accessibility

The YSA was developed in Python (v3.10.9) and is compatible with Windows and macOS. Source code and user documentation are available at [GitHub URL]. The software relies on standard Python libraries including [PyQt5, pyqtgraph, scipy, pandas, scikit-learn]. A detailed README file provides set-up instructions and usage guidelines, as well as an example data file (Lab 2025).

## Results

Using YSA’s automated detection and tracking features, we identified and visualized individual discharges within both self-limiting seizure-like events (SLSLE) and SE events. The tool enables spatial tracking of each discharge across electrodes (Figure 3B), revealing dynamic propagation patterns over time. These visualizations support detailed spatiotemporal examination of event evolution within and across brain regions.

**Figure 3:**
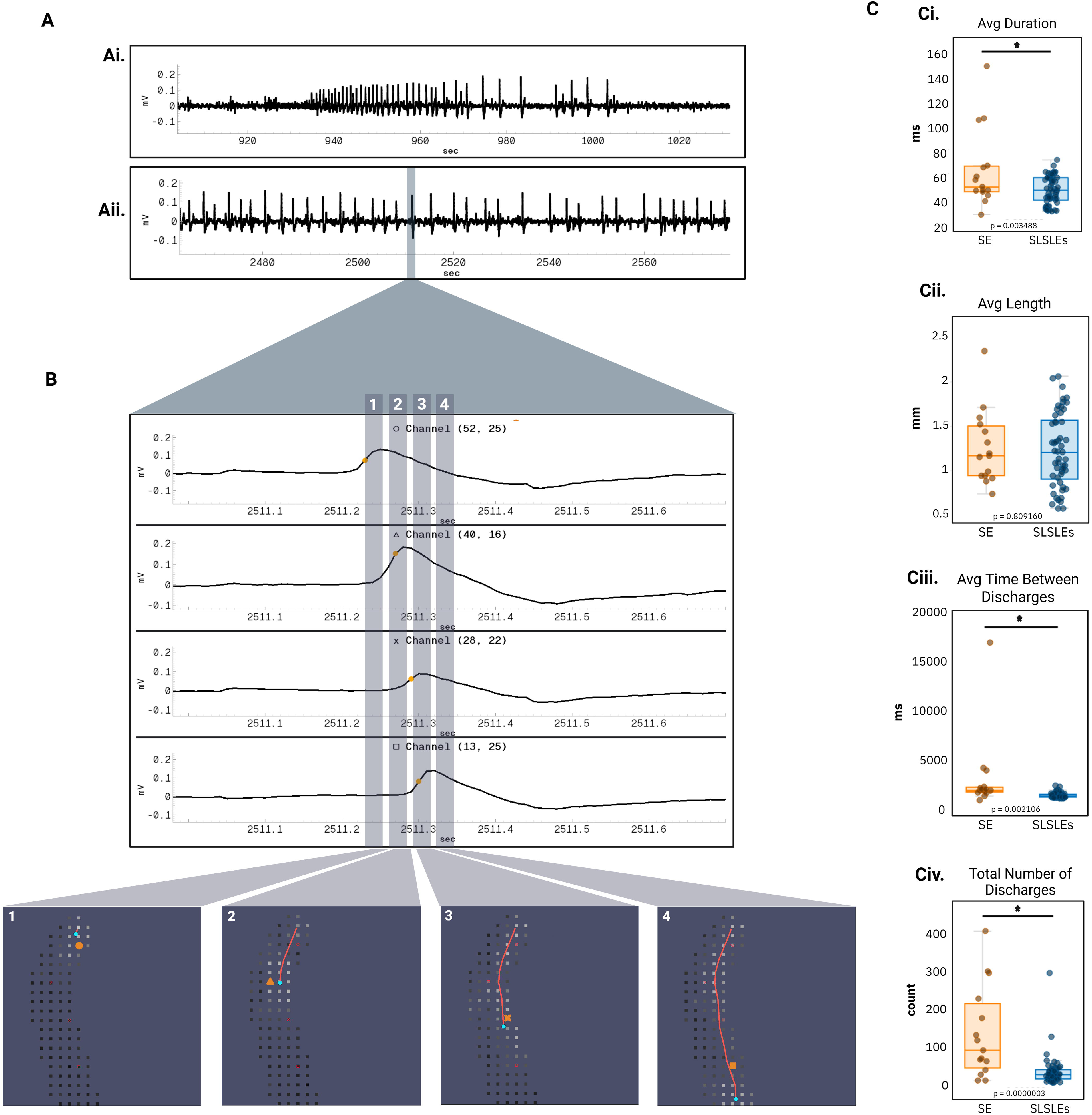
Seizure and status epilepticus discharge analysis using the YSA. **A)** Electrophysiological traces recorded from a brain slice **Ai)** Representative trace of a SLSLE**. Aii)** Representative trace of SE. **B)** Zoomed-in view of an individual discharge event. Panels 1–4 highlight distinct phases of a discharge, with corresponding electrode maps showing the spatial propagation across the slice. **C)** Quantitative comparisons between SE and SLSLE events, **Ci)** average duration (p = 0.0035), **Cii)** average path length (p = 0.81), **Ciii)** average time between discharges (p = 0.0021), and **Civ)** total average number of discharges in an SE or SLSLE event (p = 0.000003). Each scatter point represents the averaged discharge values for a single event (SE: n = 15, SLSLE: n = 53). All statistical tests were two-tailed t-tests.

**Figure 4:**
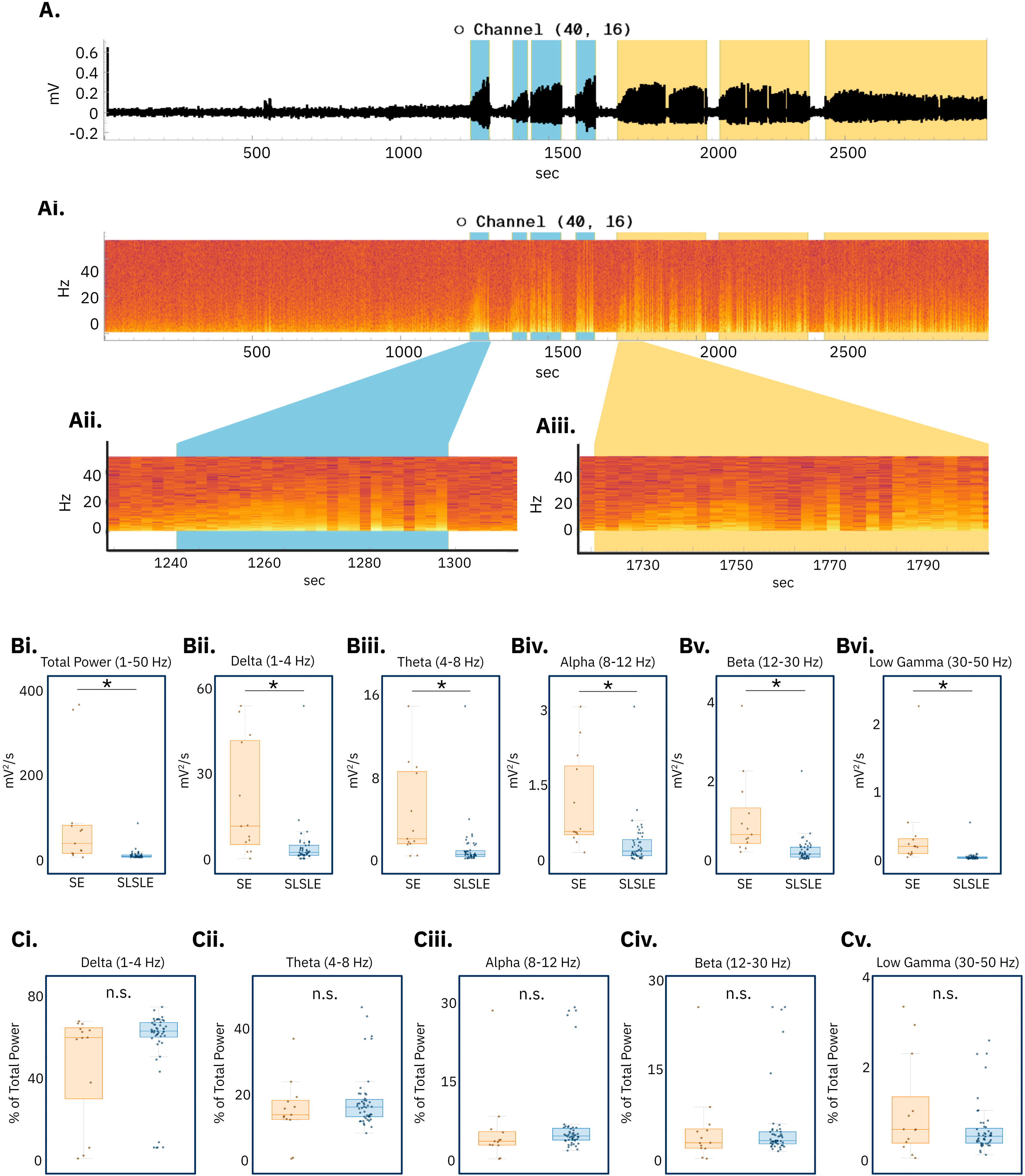
Spectral analysis of SLSLE and SE events. **A)** A tract from the YSA. **Ai.)** An example of a spectrogram generated by the YSA overlapping the selected trace. **Aii)** A zoomed in view of the spectrogram during an SLSLE. **Aiii)** A zoomed in view of the spectrogram during an SE event. **B)** The power, normalized to time, of SE events was significantly greater than the power of SLSLEs in **Bi)** total band power 1-50 Hz (p = 5.1e-5), **Bii)** delta power 1-4 Hz (p = 4.1e-5), **Biii)** theta power 4-8 Hz (p = 2.4e-4), **Biv)** alpha power 8-12 Hz (p = 8.5e-5), **Bv)** beta power 12-30 Hz (p = 1.8e-5), and **Bvi)** low gamma power 30-50 Hz (p = 5.2e-4). C) The percent of total power (normalized to time) of SE events was not significantly different than the power of SLSLEs in **Ci)** delta power 1-4 Hz (p = 0.12), **Cii)** theta power 4-8 Hz (p = 0.28), **Ciii)** alpha power 8-12 Hz (p = 0.55), **Civ)** beta power 12-30 Hz (p = 0.88), and **Cv)** low gamma power 30-50 Hz (p = 0.071). All statistical tests were two-tailed t-tests.

YSA also quantified key metrics for each detected discharge, including average duration, propagation path length, inter-discharge intervals, and total number of discharges per event (Figure 3C). As a demonstration of the tool’s analytic capabilities, we compared these features between SLSLE and SE events. SE events exhibited significantly longer discharge durations than SLSLE (Figure 3Ci). Interestingly, the average discharge length (i.e., spatial extent) did not differ significantly between states, suggesting that while SE discharges last longer in time, they do not necessarily engage in a broader spatial area (Figure 3Cii). Inter-discharge intervals were also longer in SE (Figure 3Ciii), and SE events contained a greater number of total discharges (Figure 3Civ). These results reflect and expand on differences between pharmacoresistant and pharmacosensitive seizure states and illustrate how YSA facilitates high-resolution, event-level comparisons (Barker-Haliski et al. 2017; Sanya 2004). These comparisons reveal distinct dynamic features of pharmacoresistant SE, such as increased temporal persistence without broader spatial recruitment. This insight may point to evolving mechanisms of local circuit hyperexcitability, offering clues into the network-level transitions that underline treatment-resistant seizure progression.

YSA spectral analysis has the capability to display togglable spectrograms and the ability to export spectra for events to be used in later analysis. Analysis of the spectral data demonstrated that SE events were significantly more powerful per unit time than SLSLE events both in total power (1-50 Hz, p = 5.1e-5) and across all standard frequency bands (Delta, 1-4 Hz, p = 4.1e-5; Theta, 4-8 Hz, p = 2.4e-4; Alpha, 8-12 Hz, p = 8.5e-5; Beta, 12-30 Hz, p = 1.8e-5; and Low Gamma, 30-50 Hz, p = 5.2e-4). While the SE events were more powerful, the overall composition of the power by band was not significantly different between SE and SLSLE events (Delta, 1-4 Hz, p = 0.12; Theta, 4-8 Hz, p = 0.28; Alpha, 8-12 Hz, p = 0.55; Beta, 12-30 Hz, p = 0.88; and Low Gamma, 30-50 Hz, p = 0.071). These results suggest that SE events may reflect a globally heightened state of network excitability rather than a shift in dominant frequency dynamics.

Together, these analyses demonstrate the ability of YSA to extract, visualize, and quantify both time- and frequency-domain features of HD-MEA recordings with minimal manual processing. By streamlining this process, YSA provides a robust platform for identifying subtle electrophysiological differences in seizure dynamics across experimental conditions.

## Discussion

Current MEA analysis software, while powerful, often comes with steep learning curves, limited flexibility for examining electrophysiological traces at high temporal resolution, poor adaptability to evolving recording technologies, and cost barriers (Muller et al. 2015; Obien et al. 2014; Liu et al. 2024). Many existing tools are optimized for spike and burst detection, making them less suited for tracking large-scale local field potential (LFP) activity during ictal or other network wide events (Dastgheyb, Yoo, and Haughey 2020; Hennig, Hurwitz, and Sorbaro 2019). To overcome these limitations, we developed the YSA, a user-friendly, open-access GUI that combines intuitive design with robust analytical features. YSA requires no complex setup, is compatible with future recording advancements, and is ready to use upon download. This accessibility enables researchers to perform advanced analyses, including seizure and SE dynamics in *ex vivo* recordings, without the need for specialized computational training.

While our demonstration used data from a 3Brain recording system (Blotter et al. 2024), the YSA can support data from any MEA recording system. In the supporting documentation (Lab 2025), we provide code to convert 3Brain output files (.brw) into the universal YSA format (.h5) along with data structure guidelines so users of other MEA platforms can easily reformat their recordings for use in YSA.

To showcase the utility of YSA, we applied it to a well-established acute *ex vivo* seizure model. This model provides rich, dynamic spatiotemporal data, offering an ideal test case for evaluating the tool’s analytical capabilities. While not the primary focus of this paper, our analysis illustrates how YSA enables rapid visualization, annotation, and quantification of seizure-like and SE-like events on a per-discharge basis, something often difficult to achieve with traditional software.

The YSA automatically generates raster plots and extracts electrophysiological metrics commonly used in seizure analysis (Bradley et al. 2018; Fan et al. 2019; Matsuda et al. 2022). Raster plots are essential for visualizing network-level activity, providing insights into synchronous discharges and propagation trends (Diamond et al. 2024; Wagner et al. 2015). YSA enhances this capability by integrating synchronized trace views, enabling both broad and granular analysis of seizure dynamics. Its advanced tools allow users to track individual discharges, precisely measuring duration, speed, and spatial spread across the electrode array. By enabling direct measurement of these variables, YSA helps identify distinct patterns that differentiate seizure-like states from sustained SE-like activity. These features minimize the need for custom scripting and make rigorous analysis accessible to users with varied technical backgrounds.

In addition, the YSA also supports seamless data export, simplifying integration with other analysis pipelines or inclusion in publications and presentations. By combining both standard and advanced features in a single package, YSA enhances efficiency and accessibility in HD-MEA data analysis, especially in the context of complex seizure dynamics.

In our demonstration, YSA enabled high-resolution tracking of individual discharges across the full electrode grid, revealing distinct spatiotemporal patterns associated with SE-like activity. For example, discharges during SE had significantly longer average durations than those in SLSLEs, consistent with the sustained nature of SE (Figure 3Ci) (Karunakaran, Grasse, and Moxon 2012; Treiman, Walton, and Kendrick 1990). Interestingly, the average discharge length (i.e., spatial extent) did not differ significantly between states, suggesting that while SE discharges last longer in time, they do not necessarily engage in a broader spatial area (Figure 3Cii). The average time between discharges was also significantly longer in SE, which is expected given the increased discharge duration (Figure 3Ciii). Additionally, the total number of discharges was significantly greater during SE than SLSLEs, consistent with the extended duration of SE episodes (Figure 3Civ) (Drislane et al. 2009; Johnson and Kaplan 2020; Trinka et al. 2015). Together, these findings illustrate how YSA enables detection of subtle dynamic features often missed with existing software. The ability to precisely track discharge duration, frequency, and spatial spread in real time allows researchers to distinguish between SLSLEs and more prolonged, pharmacoresistant SE-like activity. This level of granularity is critical for studies investigating mechanisms of seizure generalization, drug resistance, or network destabilization in epilepsy. By enabling high-throughput quantification of these dynamics, YSA facilitates more nuanced interpretations of network state transitions, insights that could ultimately inform both basic research and preclinical therapeutic screening.

On a broader scale, YSA’s export functions enabled spectral analysis of SLSLE and SE events, showing that while SE events exhibited significantly higher absolute power across all frequency bands, the relative power distribution remained similar. These results, again, shown as a demonstration of tool capabilities, suggest that SE events reflect a sustained and elevated level of excitability rather than a fundamentally different spectral composition (Jiruska et al. 2013; Amengual-Gual, Sanchez Fernandez, and Wainwright 2019; Holtkamp et al. 2011; Niquet et al. 2019). This kind of insight, extracted quickly from large datasets, exemplifies how YSA can accelerate data interpretation. Notably, elevated power in low-frequency bands (Delta, Theta) has been associated with impaired inhibition and disrupted network oscillations, features typical of pharmacoresistant seizure states (Avoli et al. 2016; Toutant et al. 2024; Walker 2008).

Finally, YSA successfully detected transitions from SLSLEs to SE, capturing dynamic shifts in network state that align with previous findings on SE progression (Anderson et al. 1986; Codadu et al. 2019; Burman et al. 2019; Trevelyan et al. 2023). While not intended as a biological discovery, this capacity highlights YSA’s value in identifying real-time network transitions, making it a valuable tool for researchers studying seizure generalization and pharmacoresistant SE models.

We anticipate that this platform will support future studies aimed at uncovering mechanisms of seizure evolution and drug resistance, while facilitating reproducible, high-throughput analysis of HD-MEA data (Goodchild et al. 2024; Thouta et al. 2022). Beyond seizure research, the modular and data-agnostic structure of YSA makes it broadly applicable across neuroscience and related fields. For example, the ability to visualize and quantify spatiotemporal patterns of electrical activity at the single-discharge level may prove valuable in studying network oscillations, plasticity, injury responses, or neurodevelopmental dynamics. Furthermore, because the YSA framework is customizable and compatible with a variety of HD-MEA systems, it can also be adapted for use in cardiac electrophysiology to track propagation of arrhythmic events or response to pharmacological interventions.

Importantly, by enabling researchers to rapidly extract interpretable data from large-scale recordings without specialized coding knowledge, YSA lowers the barrier to entry for electrophysiological analysis. This accessibility positions it as a valuable tool for preclinical screening and drug discovery pipelines, where efficiency and reproducibility are essential. We believe that YSA not only enhances the interpretability of complex neural datasets but also expands the experimental possibilities available to researchers across a wide range of disciplines.

